# Chemical Synthesis of Thrombostasin: An Anticoagulant Protein Produced by the Horn Fly *Haematobia irritans*

**DOI:** 10.64898/2026.09.24.754047

**Authors:** Sameer S. Kulkarni, Daniel J. Ford, Lucas Kambanis, Chenming Tang, Nisharnthi M. Duggan, Mel Shishido, Joshua W. C. Maxwell, Jorge Ripoll-Rozada, Pedro José Barbosa Pereira, Richard J. Payne

**Affiliations:** School of Chemistry, The University of Sydney, Sydney, NSW 2006, Australia; Australian Research Council Centre of Excellence for Innovations in Peptide and Protein Science, The University of Sydney, Sydney, NSW 2006, Australia; Instituto de Biomedicina y Biotecnología de Cantabria (IBBTEC), Consejo Superior de Investigaciones Científicas (CSIC)-Universidad de Cantabria, Santander, Spain; IBMC – Instituto de Biologia Molecular e Celular, Universidade do Porto, 4200-135 Porto, Portugal; i3S - Instituto de Investigação e Inovação em Saúde, Universidade do Porto, 4200-135 Porto, Portugal

## Abstract

Thrombostasin is a cysteine-free anticoagulant protein produced by the horn fly *Haematobia irritans*, featuring an unusual stretch of seven consecutive aspartate residues within the polypeptide sequence. Here, we report the first chemical synthesis of native thrombostasin and a hepta-glutamate analogue, together with assessment of their thrombin inhibitory and anticoagulant activity.

## Introduction

Cardiovascular disease (CVD) remains the leading cause of morbidity and mortality worldwide, placing immense pressure on global healthcare systems.^1-3^ A major underlying factor behind CVD is thrombosis,^4^ the obstruction of blood vessels by thrombi (clots), leading to life-threatening events such as stroke and myocardial infarction.^5^ This pathological clot formation is caused by the dysregulated activation of the coagulation cascade, culminating with the final enzyme of the pathway Factor IIa (thrombin). Thrombin catalyses the conversion of fibrinogen into insoluble fibrin that, together with platelets, makes up the thrombus. With an ageing population and increasing rates of obesity, there is predicted to be a significant rise in thrombotic events over the next decade and, while there are a number of thrombin-inhibiting anticoagulants approved for use, there are safety concerns associated with their use for specific thrombotic indications. There is therefore a growing need to identify and develop novel anticoagulant molecules for potential use in the clinic.^6^

With use beginning in medieval times, the saliva of haematophagous organisms has emerged as a rich reservoir of anticoagulants; hirudin, a 65-residue protein secreted by the medicinal leech Hirudo medicinalis, is recognised as the archetypal naturally occurring anticoagulant molecule. Other blood-feeding organisms, including mosquitoes, leeches, ticks, and flies, rely on vertebrate blood as a primary source of nutrition and have evolved to counter host haemostatic and immune responses to ensure the uninterrupted supply of a blood meal.^7^ To achieve this, these species produce a diverse repertoire of biomolecules that inhibit thrombin. At the molecular level, mature thrombin possesses a catalytic active site and two positively charged substrate-recognition regions, termed exosites I and II. The active site contains the conserved serine protease catalytic triad responsible for proteolytic cleavage of fibrinogen. Whereas exosites I and II are surface-exposed regions with basic amino acid residues, presenting strong electropositive patches at physiological pH. These exosites play a crucial role in enabling the bivalent recognition and binding of thrombin’s physiological substrates. Interestingly, most salivary anticoagulants from haematophagous organisms mimic this bivalent binding mechanism, targeting thrombin’s active site, while simultaneously engaging exosite I or exosite II to achieve exceptionally high binding affinity.^7,8^ For example, while hirudin is known to occupy the active site of thrombin, it also establishes key electrostatic interactions with exosite I via its acidic C-terminal region bearing acidic residues.^9^ Notably, a tyrosine residue (Tyr63) from the C-terminal region of hirudin has been reported to undergo post-translational sulfation, introducing an anionic sulfate group that mimics the sulfotyrosine residues found in the native fibrinogen substrate in mammals. This strategy of exploiting tyrosine sulfation to enhance thrombin inhibition through molecular mimicry extends beyond hirudin and appears to represent a broader evolutionary adaptation among natural anticoagulants from blood feeding organisms.^10^ Over the past decade, we have probed this theory through the chemical synthesis of sulfoproteins produced by leeches,^11^ mosquitoes,^12^ ticks^13^ and flies,^14^ providing access to numerous potent thrombin-inhibiting anticoagulant molecules. Moreover, we recently demonstrated that such sulfated peptides/proteins derived from different organisms can also be hybridised to generate engineered variants that exhibit ultrapotent femtomolar inhibitory activity against thrombin.^15,16^

In this work, we sought to synthesise and assess the thrombin inhibitory and anticoagulant activity of thrombostasin, a salivary protein produced by the horn fly, *Haematobia irritans*. Similar to other haematophagous organisms, horn flies must overcome the blood coagulation response of their bovine host to obtain a blood meal. In fact, horn flies are known to infest cattle in large numbers, resulting in huge economic loss due to reduced milk and beef production.^17^ In 2000, Cupp and co-workers identified salivary “factors” secreted by horn flies that inhibited thrombin activity.^18^ Subsequent studies revealed that these factors correspond to a salivary anticoagulant protein termed thrombostasin (TS), which is encoded by a polymorphic *ts* gene and facilitates blood feeding by *H. irritans*.^19^ Despite thrombin being identified as the target of TS, the kinetics of inhibition was not reported in this early work, leading to a limited understanding of its mechanism of action toward thrombin. Thrombostasin was later cloned and expressed in a baculovirus system, but the authors were unable to perform comprehensive thrombin inhibition studies due to extremely low expression yields. In our hands, we were also unable to recombinantly express TS in the bacterial host *Escherichia coli*. Herein, we report the first chemical synthesis of thrombostasin, and a structural analogue, using peptide ligation methodologies. Access to these molecules enabled assessment of their thrombin inhibitory activity, as well as potential binding modes to the protease.

## Results and discussion

The thrombostasin gene encodes a 175-residue precursor comprising an 18-residue signal peptide and a 157-residue secretory protein. However, purified thrombostasin consists of only 81 amino acids, with Ser95 as the first residue of the mature protein based on N-terminal sequencing. Consistent with this finding, Cupp and colleagues subsequently identified a proprotein convertase that cleaves the precursor between Arg94 and Ser95, generating the mature 81-residue thrombostasin.^20^ Thrombostasin does not contain any tyrosine residues and is therefore, unlike many thrombin-inhibiting anticoagulants identified from other blood feeding organisms, not post-translationally sulfated. Instead, the molecule features a unique motif containing seven consecutive negatively charged aspartate (Asp) residues along with an Asp/Glu rich region. We hypothesised that these acidic residues might be present to facilitate binding by establishing electrostatic interactions with the exosites on thrombin akin to the role of the sulfotyrosine-containing stretches found in bivalent thrombin inhibitors from other organisms.

As an 81-residue protein, and with a lack of cysteine (Cys) or alanine (Ala) residues in the N-terminal region, we chose to disconnect thrombostasin (**1**) at Met24-Asp25 and Arg55-Ala56 junctions (**Scheme 1A**). This necessitated the synthesis of three suitably functionalised peptide fragments (**2**-**4**) using Fmoc-strategy solid-phase peptide synthesis (SPPS). We envisaged fusing these fragments using kinetically controlled ligation-based assembly in the N-to-C-terminal direction using an initial rapid diselenide-selenoester ligation (DSL)^21,22^ between fragments **2** and **3**, followed by deselenisation (**Scheme 1B**). A subsequent native chemical ligation (NCL)^23^ reaction of this product with fragment **4** followed by a final desulfurisation step would then provide access to the native full-length protein. The C-terminal fragment TS(56-81) **4** was first synthesised by loading Fmoc-Ala-OH on 2-chlorotrityl chloride (2-CTC) resin, followed by peptide elongation using an automated peptide synthesiser (Scheme 2). Notably, the N-terminal Ala56 was installed as a Cys residue to facilitate NCL which could be converted back to the native Ala56 upon desulfurisation post-ligation. The synthesis of bifunctional middle fragment TS(25-55) **3** began with immobilisation of Fmoc-Arg(Pbf)-OH on hyper acid-labile 2-CTC resin, followed by elongation of the native sequence up to Asp26. A selenylated building block, Boc-(β-SePMB)Asp-OH^24^ (**5**), was then introduced at the N-terminus and the resulting peptide was cleaved from resin under mild acidic conditions. At this stage, the side-chain protected peptide was treated with 3-mercaptoethyl propionate, along with PyBOP and *i*Pr_2_NEt in DMF to introduce an alkyl thioester functionality at the C-terminus. Importantly, this step was performed at -30 °C to avoid epimerisation of the C-terminal Arg. After acidolytic cleavage, the crude PMB protected selenopeptide was subjected to PMB deprotection conditions (see ESI for details), affording the bifunctional diselenide fragment TS(25-55) **3** in 2% yield (after 62 steps from resin loading) following reverse-phase HPLC purification (**Scheme 2**). Finally, the N-terminal fragment TS(1-24) **2** bearing a C-terminal selenoester was synthesised using a side-chain anchoring strategy (Scheme 3).^25,26^ During the synthesis, we observed formation of aspartimides within the seven Asp region during SPPS, resulting in purification challenges and an extremely low isolated yield (1% over 43 steps from resin loading) for this fragment. To circumvent this deleterious side reaction, an alternate N-terminal TS(1-24)10-16E selenoester fragment **6** was synthesised wherein, the seven contiguous Asp residues were replaced with seven Glu amino acids. This enabled the generation of a thrombostasin analogue that retains the negatively charged patch in the protein but cannot form the unwanted aspartimides during chain elongation. Pleasingly, this selenoester fragment **6** was obtained in 19% yield following the same synthetic protocol (**Scheme 3**), with the improvement in synthetic yield attributed to the lack of aspartimides formed during solid-phase assembly. With the desired fragments in hand, we turned our attention to the ligation-based assembly of native full-length thrombostasin **1** and the glutamate analogue **7** (**Scheme 4**). Towards this end, the selenoester fragments [TS(1-24) **2** and **6**] were separately treated with the bifunctional diselenide fragment TS(25-55) **3** in ligation buffer (6 M Gnd•HCl, 0.1 M Na_2_HPO_4_, pH 7.0; 5 mM final concentration with respect to the monomeric selenopeptide) and the pH of resulting mixtures was carefully adjusted to 6.0. After 10 min, the additive-free DSL reactions were judged to be complete based on LC-MS analysis. Diphenyl diselenide (DPDS) generated during DSL was then extracted with Et_2_O and the ligation mixtures were treated with an equal volume of a degassed solution of 250 mM TCEP in ligation buffer to achieve complete deselenisation of β-Se moiety, which progressed to completion in 5 min. Next, the ligation products bearing a C-terminal alkyl thioester were subjected to NCL with TS(56-81) cysteinyl fragment **4**. Accordingly, the crude ligation mixtures were treated with 4 in solid form and the pH carefully adjusted to 7.0. An exogenous thiol, trifluoroethanethiol (TFET)^27^ or methyl thioglycolate (MTG) was then added to generate corresponding activated TFET or MTG thioester in situ and the reaction mixtures were incubated at 37 °C for 16 h. After completion of NCL reaction (16 h as judged by LCMS analysis, see ESI), the ligation mixtures were treated with an equal volume of a degassed solution of 500 mM TCEP and 200 mM reduced glutathione (GSH) in ligation buffer followed by a radical initiator, VA-044 (20 mM) and the reaction mixture was then incubated at 37°C for 16 h to facilitate desulfurisation of Cys56 to Ala. Subsequent HPLC purification afforded full-length native thrombostasin **1** in 20% and glutamate analogue of thrombostasin **7** in 36% isolated yield over 4 steps.

**Scheme 1.**
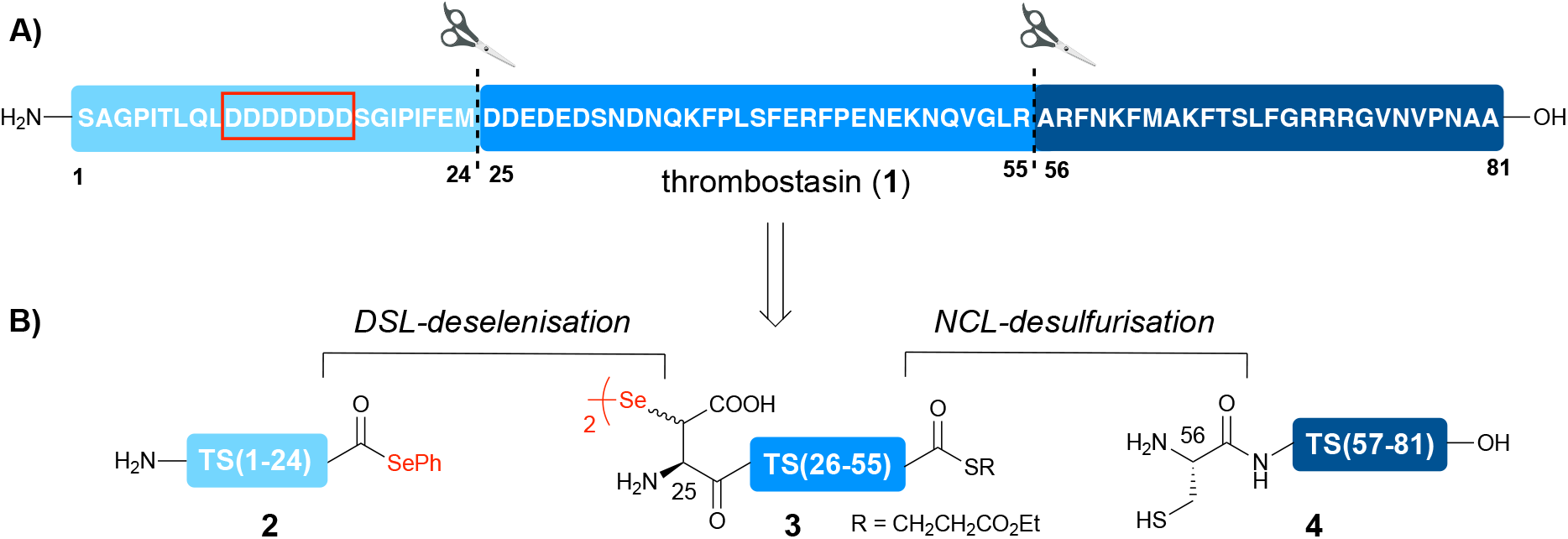
**A)** Amino acid sequence and disconnection strategy for thrombostasin. The ligation junctions are denoted with dotted lines and the red box highlights the N-terminal region featuring 7 consecutive Asp residues, **B)** Suitably functionalised required fragments for sequential ligation reactions.

**Scheme 2.**
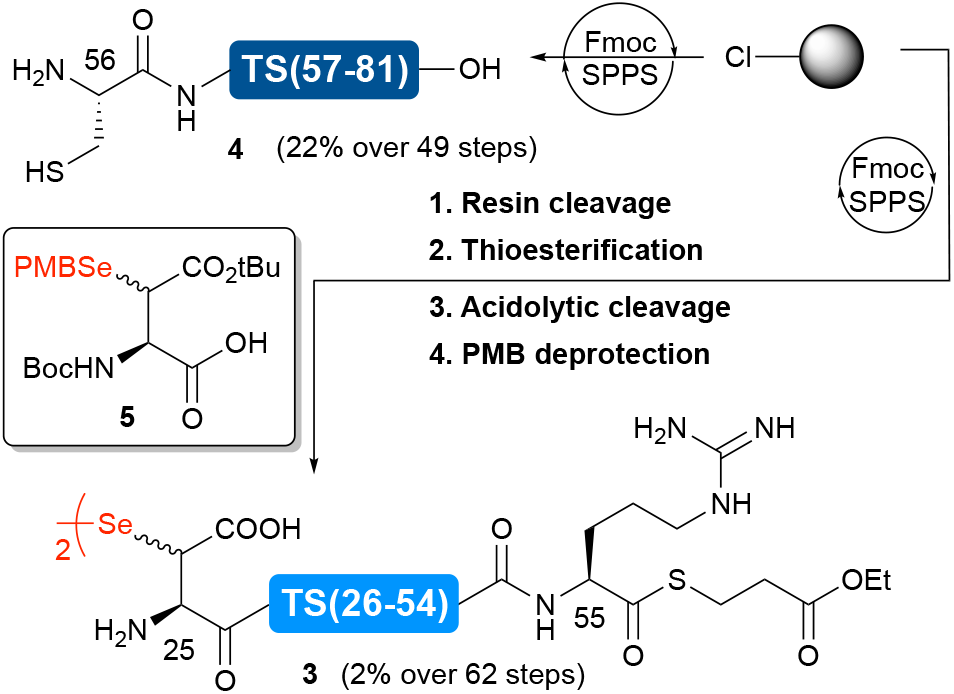
Solid-phase peptide synthesis of middle bifunctional and C-terminal fragment of thrombostasin

**Scheme 3.**
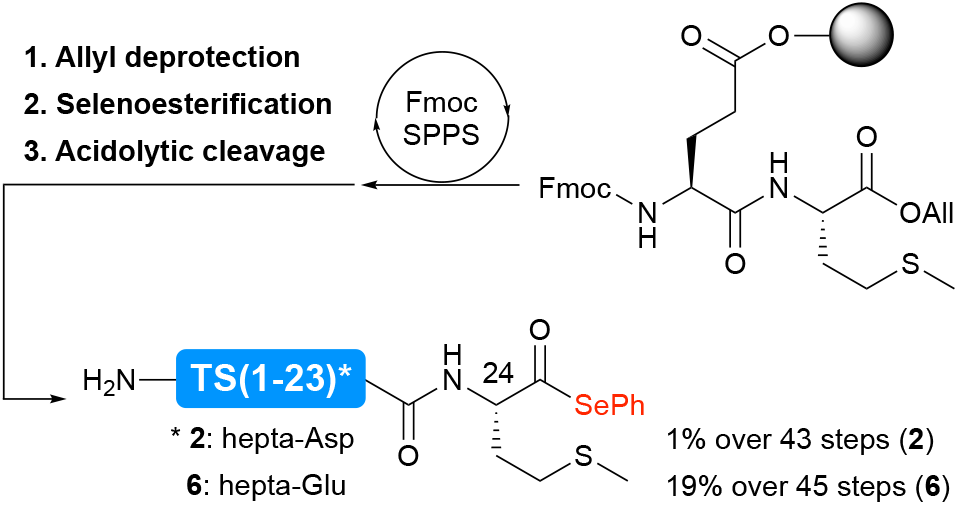
Solid-phase peptide synthesis of native and hepta-Glu variant of N-terminal fragment of thrombostasin *via* side-chain anchoring strategy

**Scheme 4.**
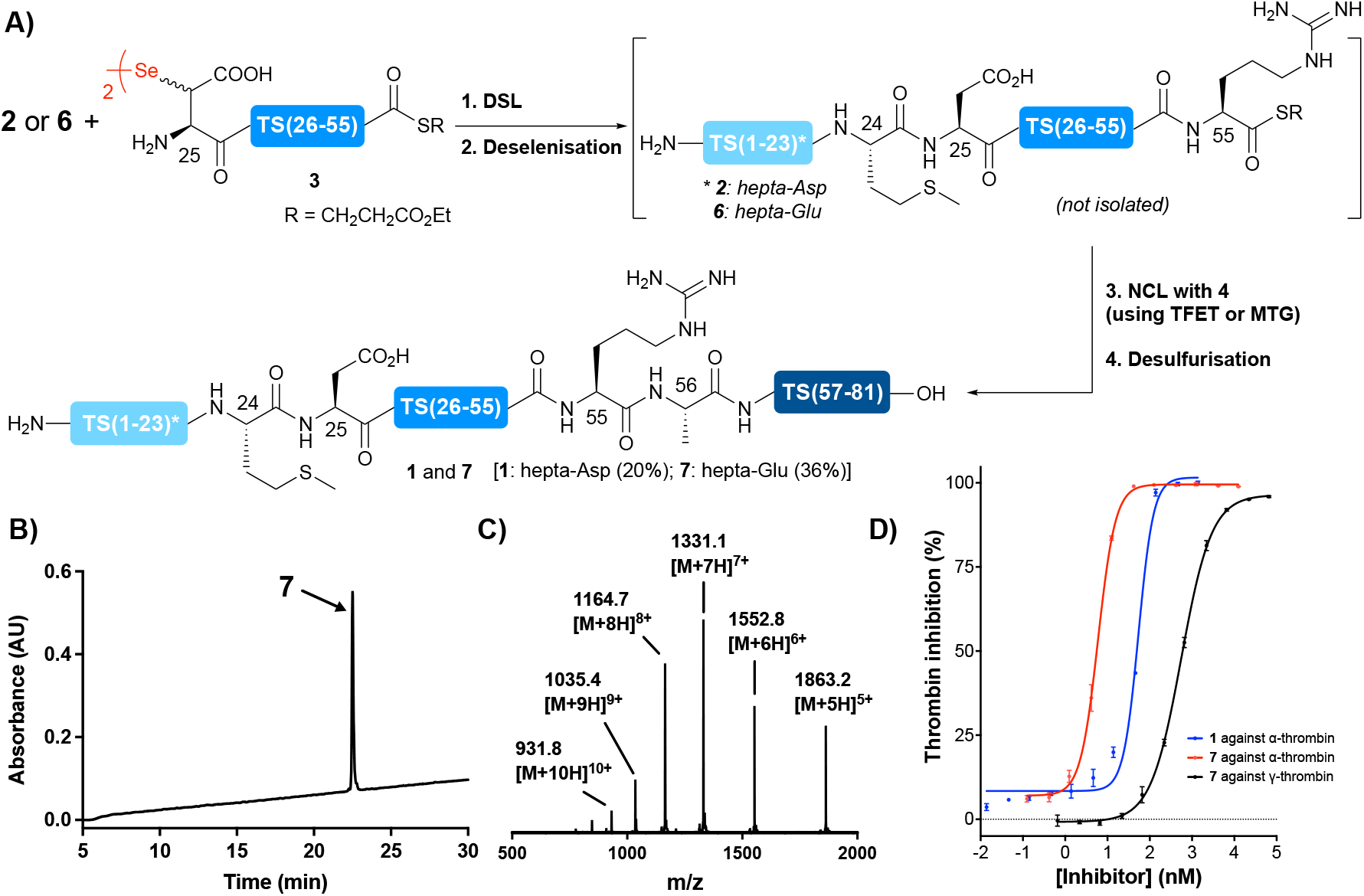
**(A)** Ligation-based assembly of the native (**1**) and hepta-Glu variant (**7**) of full-length thrombostasin *via* one-pot, sequential DSL and NCL reactions; Analytical HPLC trace **(B)** and ESI-MS **(C)** of HPLC-purified **7**; **(D)** Dose-response curves for the inhibition of α- and γ-thrombin by thrombostasin **1** and hepta-Glu variant **7**.

With native thrombostasin **1** and Glu analogue **7** in hand, we next assessed the thrombin inhibitory activity of the molecules. Specifically, a kinetic assay using Tos-Gly-Pro-Arg-p-nitroanilide as the chromogenic substrate was employed to measure the amidolytic activity of α-thrombin in the presence of varying concentrations of **1** and **7**. Interestingly, whilst native thrombostasin exhibited an IC_50_ of 54 nM, the Glu variant displayed a 10-fold improvement in inhibitory activity (IC_50_ = 6 nM). We therefore selected Glu analogue **7** for further profiling. When tested against γ-thrombin (a thrombin isoform that, despite having an impaired exosite I, still retains its amidolytic activity), the Glu variant showed ∼93-fold weaker activity compared to the activity against α-thrombin (IC_50_ = 556 nM), suggesting the involvement of exosite I in the inhibitory mode of action of thrombostasin analogue **7**.^13^ Next, Glu variant/analogue **7** was incubated with α-thrombin in Tris buffer at 37 °C to assess its stability toward its target protease. This full-length protein was cleaved at the Arg55-Ala56 junction within 1 min, suggesting that Arg55 likely occupies the P1 site of the protease when thrombostasin is bound (see ESI). To probe the importance of the C-terminal tail of thrombostasin in binding interactions with thrombin, we synthesised the truncated analogue that would result from this cleavage, [TS(1-55)10-16E]. This molecule was found to have significantly reduced inhibitory activity (IC_50_ >10 µM) against both thrombin isoforms (see ESI for details). Finally, we assessed the anticoagulant activity of hepta-Glu variant **7** using an activated Partial Thromboplastin Time (aPTT) assay (**Figure 1**). Pleasingly, thrombostasin analogue **7** showed a significant prolongation of clotting time at as little as 300 nM. Notably, the analogue showed a similar anticoagulant profile to bivalirudin albeit with slightly reduced efficacy.

**Figure 1.**
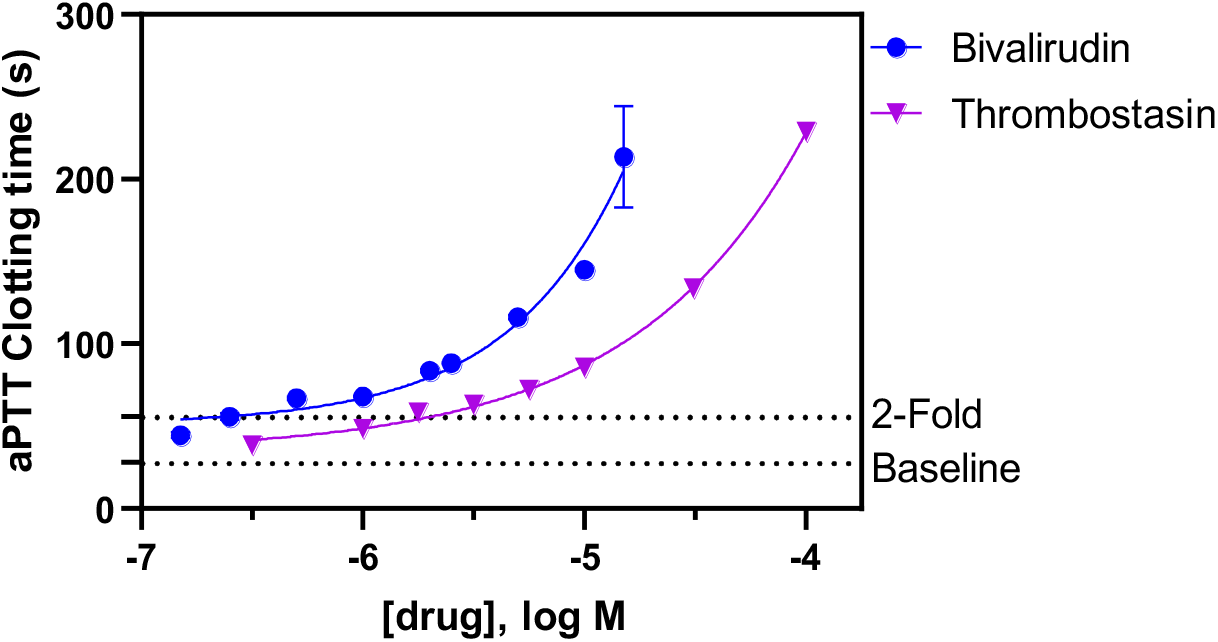
In vitro aPTT assay of thrombostasin hepta-Glu analogue **7** with bivalirudin as a positive control. Four parameter concentration–response curves for bivalirudin and thrombostasin analogue **7** were performed. Data represents the mean values ± standard error of the mean (SEM) of two independent experiments conducted in quadruplicate.

## Conclusion

In summary, we have successfully completed the first total chemical synthesis of native thrombostasin, together with an analogue with the hepta-aspartate motif found in the natural product replaced by a hepta-glutamate moiety. Synthesis was achieved through a kinetically controlled ligation of three fragments utilising consecutive DSL and NCL reactions. Both native thrombostasin and the glutamate analogue exhibited nanomolar inhibitory activity against α-thrombin, however, the glutamate analogue was ca. 10-fold more active. We also demonstrated that thrombostasin is most likely a bivalent inhibitor that occupies the active site and exosite I of thrombin and that Arg55 is most likely the residue that occupies the P1 site of the protease, with proteolytic processing resulting in significant loss of binding affinity of the resulting fragment. Finally, we demonstrated that the hepta-Glu analogue of thrombostasin possesses promising anticoagulant activity in a clinically relevant aPTT assay. Future work in our laboratories will involve attempting to co-crystallise thrombostasin and its Glu analogue with α-thrombin to unequivocally determine their binding mode to the protease. This will aid in engineering more potent and proteolytically stable anticoagulant molecules.

## Acknowledgements

The authors would like to acknowledge a New South Wales Cardiovascular Research Capacity Program Senior Researcher Grant (to R.J.P.) for funding this research.

## Data availability

Experimental procedures and characterisation data for and protocols for assays (thrombin, enzymatic cleavage and aPTT) are available in the Supporting Information document.

