## Supporting Information for "Chemical Synthesis of Thrombostasin: An Anticoagulant Protein Produced by the Horn Fly *Haematobia irritans*"

### TABLE OF CONTENTS

**Materials and Methods……………………………………………………………………..S2**

**Fmoc Solid-Phase Peptide Synthesis (Fmoc-SPPS)…………………………...……..S3**

**Resin Loading…………………………………………………………………………S3**

**Cleavage of Peptides from Resin…………………………………………………..….S4**

**Solid-phase Allyl Deprotection and Selenoesterification…………………………..……S4**

**Solution-phase Thioesterification ………………………………………………...…..S5**

**General Procedure for Additive-free Diselenide-Selenoester Ligation (DSL)……….….S5**

**Synthesis of Fmoc-Glu(OH)-Met-OAll Dipeptide…………………………………...…S6**

**Synthesis of Peptide Fragments………………………………………………………S8**

**One-pot Assembly of Native Thrombostasin (1)……………………………..………..S12**

**One-pot Assembly of Hepta-Glu Variant of Thrombostasin (7)……………..……….S14**

**Synthesis of C-terminally Truncated Thrombostasin-Glu Variant (8)………………....S16**

**Thrombin Inhibition Assay……………………………………………………………..…S17**

**Enzymatic Cleavage of Thrombostasin…………………………………………………..S18**

**activated Partial Thromboplastin Time (aPTT) assay……………………………………..S19**

**References……………………………………………………………………………S20**

### Materials and Methods

**Materials:** Reagents were used as received unless otherwise noted. Resins, coupling reagents, and amino acids were all obtained from GL Biochem, Mimotopes, or Novabiochem. *N*,*N*-dimethylformamide (DMF) was obtained as peptide synthesis grade from Merck or Labscan. All Gnd•HCl used to prepare buffers was dried *in vacuo* before use. Boc-(β-SePMB)Asp-OH was synthesised following a previously reported procedure.[1]

**Analytical HPLC** was performed on a Waters Alliance e2695 HPLC system equipped with a 2998 PDA detector (λ = 210-400 nm). A Waters XBridge C8 or C18 BEH, 130Å, 3.5 µm, 4.6 mm × 150 mm column was used with a flow rate of either 0.4 mL/min or 1 mL/min and a mobile phase composed of water (solvent A) and MeCN (solvent B) with 0.1 vol.% trifluoroacetic acid (TFA) as an additive.

**Analytical UPLC** was performed on a Waters Acquity UPLC system equipped with a Sample Manager FTN, Quaternary Solvent Manager (H-Class) and a PDA eλ detector (λ = 210-400 nm) with chromatograms extracted at λ = 214 nm. A Waters Acquity C8 or C18 BEH, 130Å, 1.7 µm, 2.1 mm × 50 mm column was used with a flow rate of 0.6 mL/min and a mobile phase composed of water (solvent A) and MeCN (solvent B) with 0.1 vol.% TFA as an additive.

The analysis of all analytical HPLC and UPLC chromatograms was conducted using Empower 3 Pro software (2010).

**Preparative HPLC:** Preparative reverse-phase HPLC was performed using a Waters 600 Multisolvent Delivery System and Waters 500 pump with 2996 photodiode array detector or Waters 490E Programmable Wavelength Detector operating at 214 and 280 nm. All peptide fragments were purified using a Waters XBridge preparative column BEH, 130 Å, 5 µm, 40 mm × 160 mm. All ligation products and full-length proteins were purified using a Waters XBridge BEH, 130 Å, 5 µm, 40 mm × 160 mm C8 preparative column.

**UPLC-MS:** UPLC-MS was performed on a Shimadzu LC-MS 2020 system equipped with a Nexera X2 LC-30AD pump and a Nexera X2 SPD-M30A diode array detector coupled to a Shimadzu 2020 electron spray ionization-mass spectrometer (ESI-MS) operating in positive mode. Peptides and proteins were analysed on a C18 (XBridge BEH C18, 130Å, 1.7 µm, 2.1 mm × 50 mm and C8 (XBridge BEH C8, 1.7 µm, 2.1 mm × 50 mm) UPLC column respectively, using a gradient of 0-70% B over 5 min, 0.1 vol.% formic acid (FA). All mass spectra extracted from UPLC-MS data are from the total ion count of the entire UPLC gradient.

**HRMS-ESI:** HRMS-ESI mass spectra were recorded with a Bruker SolariX 2xR 7T Fourier Transform Ion Cyclotron Resonance Mass Spectrometer (FTICR) controlled by ftmsControl version 2.3.0. ESI samples were administered by direct syringe infusion from water/acetonitrile (1:1 v/v) containing 0.1 vol.% FA. Q1 mass was set to the presented m/z signal and continuous accumulation of selected ions (CASI) quadrupole isolation applied with a width of 10-50 m/z.

**Fmoc Solid-Phase Peptide Synthesis (Fmoc-SPPS)**

**Resin Loading**

**Loading 2-chlorotrityl chloride resin:** 2-chlorotrityl chloride resin (1.22 mmol/g loading) was swollen in $CH_2Cl_2$ for 30 min. A solution of Fmoc protected amino acid (3 equiv. relative to theoretical loading) and *i*$Pr_2NEt$ (12 equiv.) in $CH_2Cl_2$ was added to the resin and agitated for 16 hours at rt. The loading solution was drained and the resin was washed with DMF (5 × 3 mL), $CH_2Cl_2$ (5 × 3 mL), and DMF (5 × 3 mL). The resin was then treated with a capping solution of $CH_2Cl_2$/MeOH/ *i*$Pr_2NEt$ (17:2:1 v/v/v, 4 mL) for 1 h and subsequently washed with DMF (5 × 3 mL), $CH_2Cl_2$ (5 × 3 mL), and DMF (5 × 3 mL).

**Quantification of resin loading:** A measured quantity of approximately 10 mg of the loaded resin was treated with a solution of 2 vol.% 1,8-diazabicyclo[5.4.0]undec-7-ene (DBU) in DMF (2 mL) and agitated for 30 min. The solution was removed from resin and diluted to 10 mL with MeCN. A 1 mL aliquot of this resultant solution was taken up and diluted to 12.5 mL with MeCN. The absorbance of the DBU-fulvene adduct ($\lambda$ = 304 nm, $\varepsilon$ = 9254 $M^{-1}$ $cm^{-1}$) was measured to estimate the resin loading.

**General method using SYRO I Automated Synthesizer:** Fmoc-SPPS was performed on a Biotage SYRO I automated synthesizer on a 50 µmol scale. Solutions of Fmoc-AA-OH (0.5

M) and Oxyma (0.55 M) in DMF, and *N,N'*-di*iso*propylcarbodiimide (DIC, 0.5 M) in DMF were prepared for coupling. General synthetic steps were:

*Fmoc deprotection*: The resin-bound peptide was shaken in a solution of 40 vol.% piperidine in DMF (800 μL, 4 min). The deprotection solution was then drained and the resin was treated again with a solution of 20 vol.% piperidine in DMF (800 μL, 4 min). The deprotection solution was then drained and the resin was washed with DMF (4 × 1.0 mL).

*Coupling*: The resin-bound peptide was shaken in a solution of Fmoc-AA-OH (4 equiv., 0.17 M), Oxyma (4.4 equiv., 0.18 M) and DIC (4 equiv., 0.17 M) in DMF for 45 min at 40 °C. The coupling solution was drained, and the resin was treated with a fresh coupling solution for 45 min at 40 °C and then washed with DMF (4 × 800 μL).

*Capping*: The resin-bound peptide was shaken in a solution of 5 vol.% $Ac_2O$ and 10 vol.% $i Pr_2NEt$ in DMF (800 μL, 6 min). The capping solution was then drained, and the resin was washed with DMF (4 × 800 μL).

#### Cleavage of Peptides from Resin

**Acidolytic cleavage from resin along with deprotection of side-chain protecting groups:** The resin (50 μmol) was first washed with $CH_2Cl_2$ (2 × 3 mL), dried and treated with a mixture of trifluoroacetic acid (TFA), $i Pr_3SiH$ (TIS) and water (18:1:1 v/v/v, 5 mL) with gentle agitation at rt for 2 h. The resin was then filtered and washed with TFA (2 × 3 mL) and the combined filtrates were concentrated under a stream of $N_2$. $Et_2O$ (40 mL) was added and the suspension cooled to 0 °C for 10 min. The precipitate was pelleted by centrifugation at 6800 rcf for 10 min at 0 °C and the supernatant decanted.

#### Solid-phase Allyl Deprotection and Selenoesterification

The resin (25 μmol) was swollen in dry $CH_2Cl_2$ for 30 min and after draining the solvent, the resin was treated with a solution of $Pd(PPh_3)_4$ (1 equiv.) and $PhSiH_3$ (40 equiv.) in dry $CH_2Cl_2$ (2.5 mL). The resin was shaken for 1 h and the procedure was repeated to achieve complete allyl deprotection. Afterwards, the resin was washed with $CH_2Cl_2$ (5 × 3 mL), DMF (5 × 3 mL), $CH_2Cl_2$ (5 × 3 mL) and treated with a solution of DPDS (30 equiv.) and $n Bu_3P$ (30 equiv.) in dry DMF (2.5 mL) at 0 °C for 3 h, resulting in C-terminal selenoesterification. The resin was then washed with $CH_2Cl_2$ (5 × 3 mL), DMF (5 × 3 mL), $CH_2Cl_2$ (5 × 3 mL) and subjected to acidolytic cleavage followed by RP-HPLC purification.

**Solution-phase Thioesterification**

The peptide thioester was prepared on a hyper acid-labile 2-CTC resin using the standard Fmoc-strategy SPPS method described above. After elongation of the target amino acid sequence, the peptide was cleaved from resin using 30 vol.% 1,1,1,3,3,3-hexafluoroisopropanol (HFIP) in $CH_2Cl_2$ (3 mL for 25 μmol, 1.5 h), leaving the side-chain protecting groups intact. After concentrating the solution, the crude peptide was subjected to thioesterification using ethyl 3-mercaptopropionate (30 equiv.), $i$$Pr_2NEt$ (5 equiv.) and PyBOP (benzotriazol-1-yloxytripyrrolidinophosphonium hexafluorophosphate, 5 equiv.) in DMF at -30 °C for 3 h. It is crucial to perform this step at -30 °C to avoid epimerization of the C-terminal amino acid. The reaction mixture was then concentrated using a stream of $N_2$ and subjected to acidolytic cleavage followed by RP-HPLC purification.

**General Procedure for Additive-free Diselenide-Selenoester Ligation (DSL)**

Peptide selenoester (1.2-1.4 equiv.) and the peptide diselenide dimer (1 equiv. with respect to the monomer) were dissolved separately in ligation buffer (6 M Gnd•HCl, 100 mM $Na_2HPO_4$, pH 7.2) to a concentration of 2.5 mM with respect to the peptide diselenide dimer (5 mM with respect to the monomer). The solutions were combined in one portion and the pH was carefully adjusted to 6.0 with 1 M NaOH solution. The ligation was allowed to proceed at room temperature and monitored by UPLC-MS to ensure complete consumption of the selenopeptide fragment.

**Synthesis of Fmoc-Glu(OH)-Met-OAll Dipeptide:**

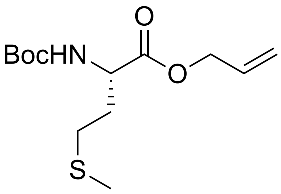


Boc-Met-OH (2 g, 8.0 mmol, 1 equiv.) was dissolved in DMF (20 mL) and treated with allyl bromide (0.76 mL, 8.8 mmol, 1.1 equiv.) and *i*$Pr_2NEt$ (4.2 mL, 24 mmol, 3 equiv.) for 16 h at room temperature. The reaction mixture was then diluted with water and extracted with EtOAc. The combined organic layers were washed with 1 M HCl (×3) and $H_2O$ (×3), dried over $Na_2SO_4$ and concentrated *in vacuo* to afford Boc-Met-OAll (2.32 g, 8.0 mmol, quantitative conversion), which was taken to the next step without purification.

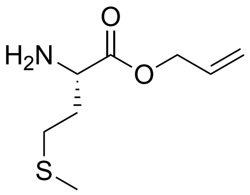


Boc-Met-OAll was dissolved in 1:1 (v/v) TFA/$CH_2Cl_2$ (20 mL) and stirred at rt for 1 h to achieve complete Boc deprotection. The reaction mixture was then concentrated in *vacuo* and residual TFA was azeotroped using toluene (×3) to afford $H_2N$-Met-OAll (1.07 g, 4.8 mmol, 61%), which was taken to the next step without purification.

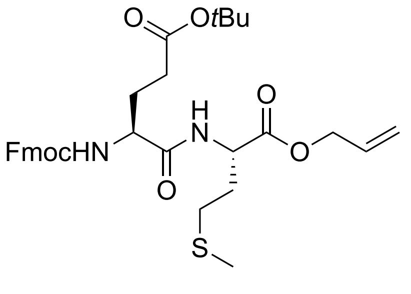


Fmoc-Glu(O*t*Bu)-OH (2.55 mg, 6.0 mmol, 1.2 equiv.) and HATU (2.26 mg, 5.95 mmol, 1.19 equiv.) were dissolved in DMF (30 mL), treated with NMM (3.2 mL, 24 mmol, 4.8 equiv.) and the resulting solution was added to H-Met-OAll (945 mg, 5.0 mmol, 1 equiv.). After stirring for 2 h at rt, the reaction mixture was concentrated under a gentle stream of $N_2$, diluted with $CH_2Cl_2$ (100 mL) and washed with water (80 mL), 2 M HCl (80 mL), sat. $NaHCO_3$ (80 mL) and brine (100 mL). The organic layer was dried over $MgSO_4$ and concentrated *in vacuo*. The crude residue was purified via silica flash chromatography (2:8 EtOAc/petroleum benzine) to afford Fmoc-Glu(O*t*Bu)-Met-OAll as a white solid (2.02 g, 3.4 mmol, 68%). **$^1$H NMR** ($CDCl_3$, 400 MHz) δ (ppm): 7.76 (dd, *J* = 7.6, 1.1 Hz, 2H, *Fmoc-Ar-**H***), 7.59 (d, *J* = 7.5 Hz, 2H, *Fmoc-Ar-**H***), 7.43 – 7.36 (m, 2H, *Fmoc-Ar-**H***), 7.31 (td, *J* = 7.4, 1.2 Hz, 2H, *Fmoc-Ar-**H***), 7.06 (s, 1H, ***NH**-amide*), 5.90 (tt, *J* = 11.0, 5.4 Hz, 1H, *All-CH2**CH**CH2*), 5.74 (s, 1H, Fmoc-***NH***), 5.33 (d, *J* = 17.1 Hz, 1H, *All-CH2CHC**H2A***), 5.26 (d, *J* = 10.5 Hz, 1H, *All-CH2CHC**H2B***), 4.76 – 4.68 (m, 1H, *Met-**αH***), 4.64 (d, *J* = 5.8 Hz, 2H, *All-**CH2**CHCH2*), 4.38 (d, *J* = 7.1 Hz, 2H, *Fmoc-**CH2***), 4.31 – 4.25 (m, 1H, *Glu-**αH***), 4.21 (t, *J* = 7.1 Hz, 1H, *Fmoc-C**H***), 2.53 (t, *J* = 7.4 Hz, 2H, *Met-**γH2***), 2.42 (t, *J* = 8.2 Hz, 2H, *Glu-**γH2***), 2.18 (dt, *J* = 13.2, 6.9 Hz, 2H, *Met-**βH2***), 2.07 (s, 3H), 1.98 (dt, *J* = 14.1, 7.1 Hz, 2H, *Glu-**βH2***), 1.47 (s, 9H, *t*Bu). **$^{13}$C NMR** ($CDCl_3$, 100 MHz) δ (ppm): 173.1, 171.3, 171.2, 156.2, 143.9, 141.3, 131.4, 127.7, 127.1, 125.1, 120.0, 119.1, 81.2, 67.2, 66.2, 54.2, 51.8, 47.2, 31.7, 31.5, 30.0, 28.4, 28.1, 15.5. **HRMS**: (ESI+) Calc. for 619.24484 $[M+Na]^+$, Found: 619.24533. **IR** (ATR): $v_{max}$ = 3295, 3068, 2976, 2919, 1731, 1687, 1648, 1532, 1278, 1151, 737 $cm^{-1}$.

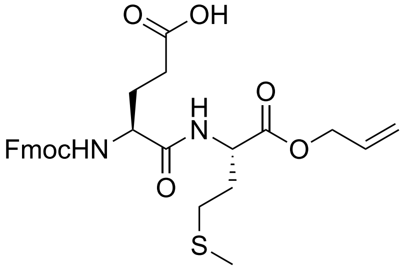


Fmoc-Glu(O*t*Bu)-Met-OAll dipeptide (895 mg, 1.5 mmol) was dissolved in 1:1 (v/v) TFA/$CH_2Cl_2$ (20 mL) and stirred for 1 h at rt to achieve complete deprotection of the *tert*-butyl ester moiety. The reaction mixture was then concentrated *in vacuo* and residual TFA was azeotroped using toluene (×3) to afford Fmoc-Glu(OH)-Met-OAll as an off-white solid (784 mg, 1.45 mmol, 97%). **$^1$H NMR** ($CDCl_3$, 400 MHz) δ (ppm): 7.73 (dd, *J* = 7.6, 1.1 Hz, 2H, *Fmoc-Ar-**H***), 7.56 (d, *J* = 7.5 Hz, 2H, *Fmoc-Ar-**H***), 7.45 (d, *J* = 7.9 Hz, 1H, ***NH**-amide*), 7.40 – 7.34 (m, 2H, *Fmoc-Ar-**H***), 7.31 – 7.25 (m, 2H, *Fmoc-Ar-**H***), 6.10 (d, *J* = 8.6 Hz, 1H, ***NH***-Fmoc), 5.93 – 5.81 (m, 1H, *All-CH2**CH**CH2*), 5.30 (d, *J* = 17.2 Hz, 1H, *All-CH2CHC**H2A***), 5.23 (d, *J* = 10.5 Hz, 1H, *All-CH2CHC**H2B***), 4.76 – 4.66 (m, 1H, *Met-**αH***), 4.60 (d, *J* = 4.6 Hz, 2H, *All-**CH2**CHCH2*), 4.54 – 4.46 (m, 1H, *Glu-**αH***), 4.40 – 4.28 (m, 2H, *Fmoc-**CH2***), 4.18 (t, *J* = 7.2 Hz, 1H, *Fmoc-C**H***), 2.53 – 2.46 (m, 4H, *Glu-**γH2***, *Met-**γH2***), 2.14 (dt, *J* = 13.4, 7.0 Hz, 2H, *Met-**βH2***), 2.01 (s, 3H, *Met-SC**H3***), 1.98 (t, *J* = 7.0 Hz, 2H, *Glu-**βH2***). **$^{13}$C NMR** ($CDCl_3$, 100 MHz) δ (ppm): 176.4, 171.9, 171.3, 156.6, 143.7, 141.3, 131.4, 127.8, 127.1, 125.1, 120.0,

119.1, 67.5, 66.2, 53.7, 51.8, 47.0, 31.2, 30.0, 29.7, 28.1, 15.4. **HRMS**: (ESI+) Calc. for 563.18224 $[M+Na]^+$, Found: 563.18230. **IR** (ATR): $\nu_{max}$ = 3295, 3067, 3007, 2915, 1732, 1688, 1649, 1532, 1368, 1278, 1155, 736 $cm^{-1}$.

**Synthesis of Peptide Fragments:**

*TS(1-24)-SePh (**2**)*

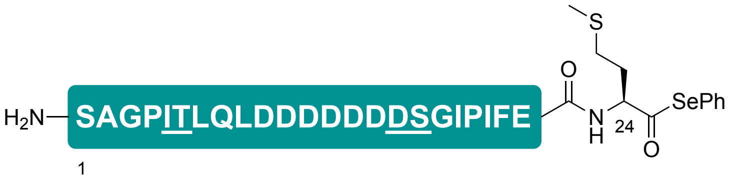


The Fmoc-Glu(OH)-Met-OAll dipeptide was loaded onto 2-CTC resin via side-chain carboxylate of Glu (140 µmol) and the peptide sequence was elongated using standard Fmoc-SPPS protocol described in the general methods section on an automated peptide synthesiser (Syro I). The underlined residues were incorporated as the corresponding pseudoproline dipeptides (DS = Fmoc-Asp(O*t*Bu)-Ser($\Psi^{Me,Me}$Pro)-OH, IT = Fmoc-Ile-Thr($\Psi^{Me,Me}$Pro)-OH) and Fmoc deprotections were carried out using 20% piperidine in DMF with 0.1 M Oxyma after coupling DS pseudoproline to suppress aspartimide formation. The fully elongated resin-bound peptide was then subjected to allyl deprotection followed by selenoesterification using conditions described in the general methods section. Finally, the peptide selenoester was treated with 85:5:5:5 (v/v/v/v) TFA/*i*$Pr_3SiH$/$H_2O$/thioanisole for 2 h. After filtering off the resin, the filtrate was concentrated under a stream of nitrogen, precipitated using cold $Et_2O$ and centrifuged. The resulting crude peptide pellet was dissolved in ligation buffer (6 M Gnd•HCl, 100 mM $Na_2HPO_4$, pH 5) and subjected to preparative RP-HPLC purification (0 to 20% B over 3 min then 20 to 80% B over 60 min, 0.1 vol.% TFA), affording **2** as a fluffy white solid after lyophilisation (4.5 mg, 1% over 43 steps).

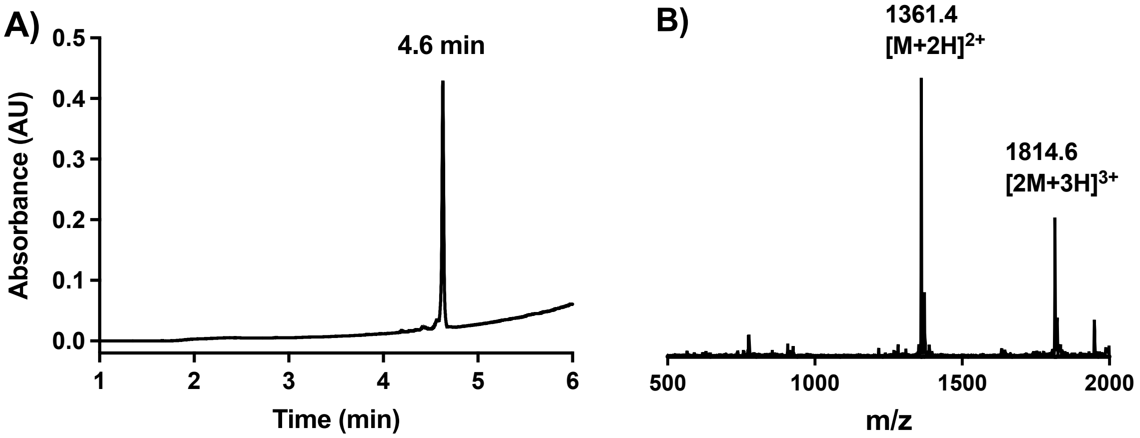

**Figure S1. A)** Analytical UPLC trace of RP-HPLC purified TS(1-24)-SePh (**2**), $R_t$ = 4.6 min (0% B for 1 min, then 0 to 80% B over 5 min, 0.1 vol.% TFA, λ = 230 nm); **B)** ESI-MS of **2**, Expected Mass (ESI+): 1360.4 $[M+2H]^{2+}$, 1813.5 $[2M+3H]^{3+}$. Mass Found (ESI+): 1361.4 $[M+2H]^{2+}$, 1814.6 $[2M+3H]^{3+}$.

*TS(1-24)10-16E-SePh (**6**)*

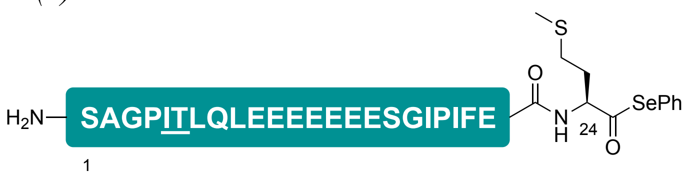


This peptide selenoester fragment TS(1-24)E-SePh was synthesised on 50 µmol scale using a similar synthetic protocol as described for TS(1-24)-SePh. In this case, only one pseudoproline dipeptide (Fmoc-Ile-Thr($\Psi^{Me,Me}$Pro)-OH) was incorporated and standard Fmoc deprotection conditions (20% piperidine in DMF) were employed throughout the entire peptide sequence. The crude peptide was purified using RP-HPLC (1 to 60% B over 50 min, 0.1 vol.% TFA) affording **6** as a white fluffy solid after lyophilisation (27 mg, 19% over 45 steps).

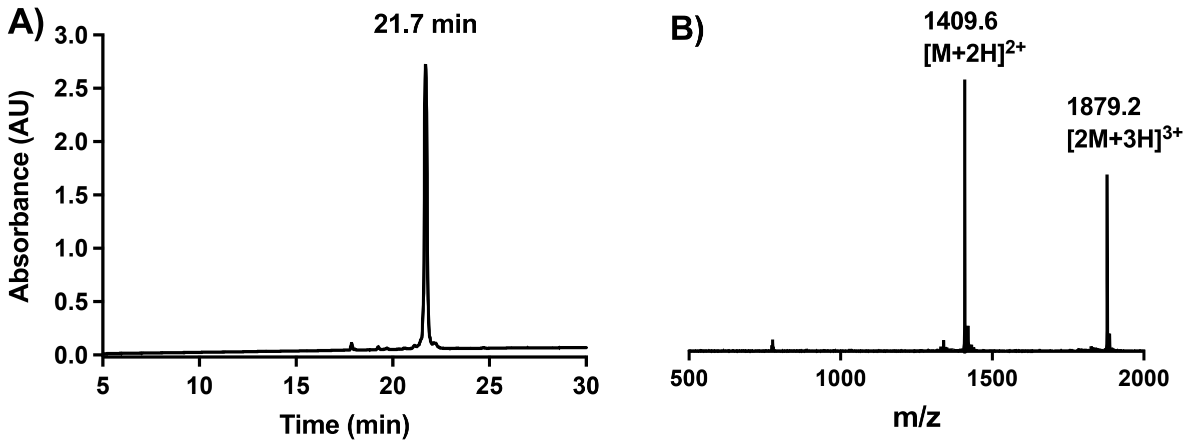


**Figure S2. A)** Analytical HPLC trace of RP-HPLC purified TS(1-24)10-16E-SePh (**6**), $R_t$ = 21.7 min (1 to 70% B over 30 min, 0.1 vol.% TFA, λ = 214 nm); **B)** ESI-MS of **6**, Expected Mass (ESI+): 1409.5 $[M+2H]^{2+}$, 1879.0 $[2M+3H]^{3+}$. Mass Found (ESI+): 1409.6 $[M+2H]^{2+}$, 1879.2 $[2M+3H]^{3+}$.

*Bifunctional TS(25-55) prior to PMB deprotection (**3S**)*

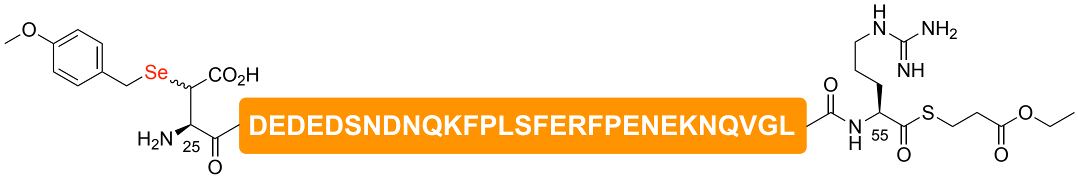


Fmoc-Arg(Pbf)-OH was loaded onto 2-CTC resin (200 µmol) and the peptide sequence was elongated up to Asp26 using the standard Fmoc-SPPS protocol described in the general methods section on an automated peptide synthesiser (Syro I). The resin was split at this point and Boc-(β-SePMB)Asp-OH building block was coupled at the N-terminus to a portion of resin (50 µmol) using standard DIC/Oxyma coupling conditions. The peptide was then subjected to HFIP cleavage followed by solution-phase thioesterification as described in the general methods section. The reaction mixture was then concentrated using a stream of $N_2$ and subjected to acidolytic cleavage [TFA/*i*$Pr_3SiH/H_2O$ (90:5:5)] for 2 h. Upon RP-HPLC purification (1 to 50% B over 50 min, 0.1 vol.% TFA), the peptide thioester bearing N-terminal PMB protected β-Se Asp (**3S**) was isolated as a white fluffy solid after lyophilisation (20.4 mg, 10% over 60 steps).

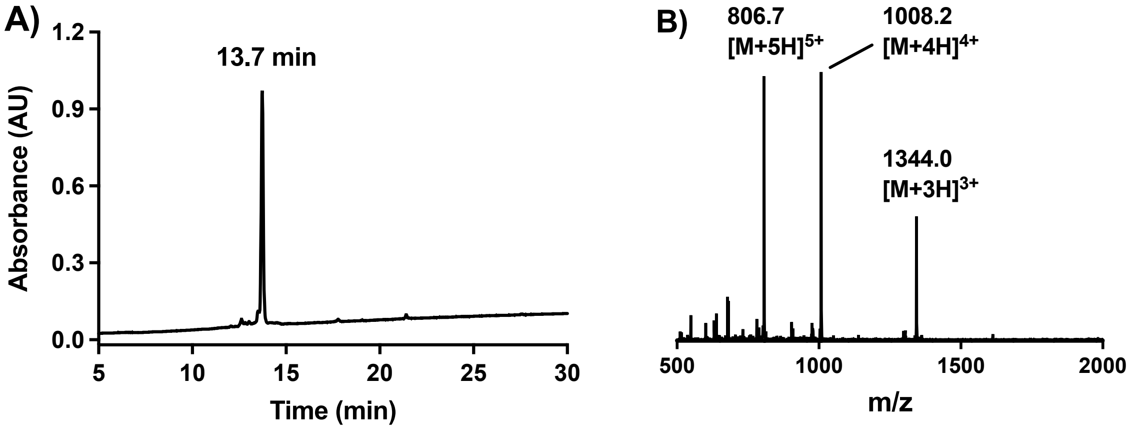


**Figure S3. A)** Analytical HPLC trace of RP-HPLC purified TS(25-55) prior to PMB deprotection (**3S**), $R_t$ = 13.7 min (1 to 60% B over 30 min, 0.1 vol.% TFA, λ = 214 nm); **B)** ESI-MS of **3S**, Expected Mass (ESI+): 1343.7 $[M+3H]^{3+}$, 1008.1 $[M+4H]^{4+}$, 806.6 $[M+5H]^{5+}$. Mass Found (ESI+): 1344.0 $[M+3H]^{3+}$, 1008.2 $[M+4H]^{4+}$, 806.7 $[M+5H]^{5+}$.

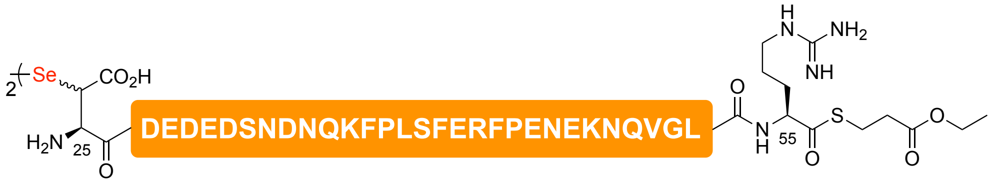


**PMB deprotection:** The PMB protected TS(25-55) alkyl thioester fragment was dissolved in a solution of TFA/DMSO/6 M Gnd•HCl, 1 M HEPES buffer (3:1:1) and the resulting mixture was incubated at room temperature for 30 min. Upon complete PMB deprotection, the reaction mixture was subjected to RP-HPLC purification (1 to 40% B over 40 min, 0.1 vol.% TFA), affording the desired bifunctional diselenide fragment as a white fluffy solid after lyophilisation (4.5 mg, 23% over 2 steps).

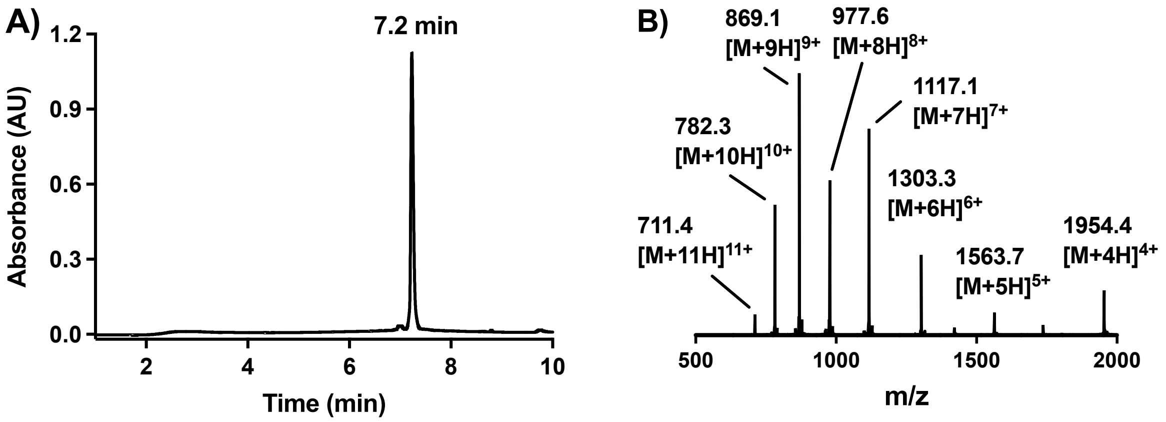


**Figure S4. A)** Analytical UPLC trace of RP-HPLC purified TS(25-55) diselenide (**3**), $R_t$ = 7.2 min (1 to 60% B over 10 min, 0.1 vol.% TFA, $\lambda$ = 214 nm); **B)** ESI-MS of **3**, Expected Mass (ESI+): 1954.5 $[M+4H]^{4+}$, 1563.8 $[M+5H]^{5+}$, 1303.3 $[M+6H]^{6+}$, 1117.3 $[M+7H]^{7+}$, 977.8 $[M+8H]^{8+}$, 869.2 $[M+9H]^{9+}$, 782.4 $[M+10H]^{10+}$, 711.4 $[M+11H]^{11+}$. Mass Found (ESI+): 1954.4 $[M+4H]^{4+}$, 1563.7 $[M+5H]^{5+}$, 1303.3 $[M+6H]^{6+}$, 1117.1 $[M+7H]^{7+}$, 977.6 $[M+8H]^{8+}$, 869.1 $[M+9H]^{9+}$, 782.3 $[M+10H]^{10+}$, 711.4 $[M+11H]^{11+}$.

*TS(56-81) (**4**)*

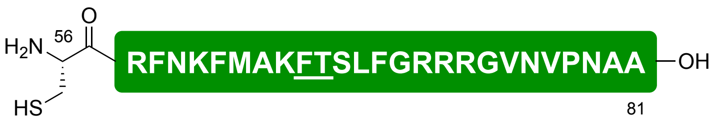


Fmoc-Ala-OH was loaded onto 2-CTC resin (50 µmol) and the peptide sequence was elongated using standard Fmoc-SPPS protocol described in the general methods section on an automated peptide synthesiser (Syro I). The underlined residues were incorporated as a pseudoproline dipeptide (FT = Fmoc-Phe-Thr($\Psi^{Me,Me}$Pro)-OH). The fully elongated peptide (50 µmol) was then treated with TFA/*i*$Pr_3SiH/H_2O$/EDT/$NH_4I$ (95:5:5:2.5:1.5) for 2 h. After filtering off the resin, the filtrate was concentrated under a stream of nitrogen, precipitated using cold $Et_2O$ and centrifuged. The resulting crude peptide was purified using RP-HPLC (1 to 40% B over 40 min, 0.1 vol.% TFA), affording **4** as a white fluffy solid after lyophilisation (33 mg, 22% over 49 steps).

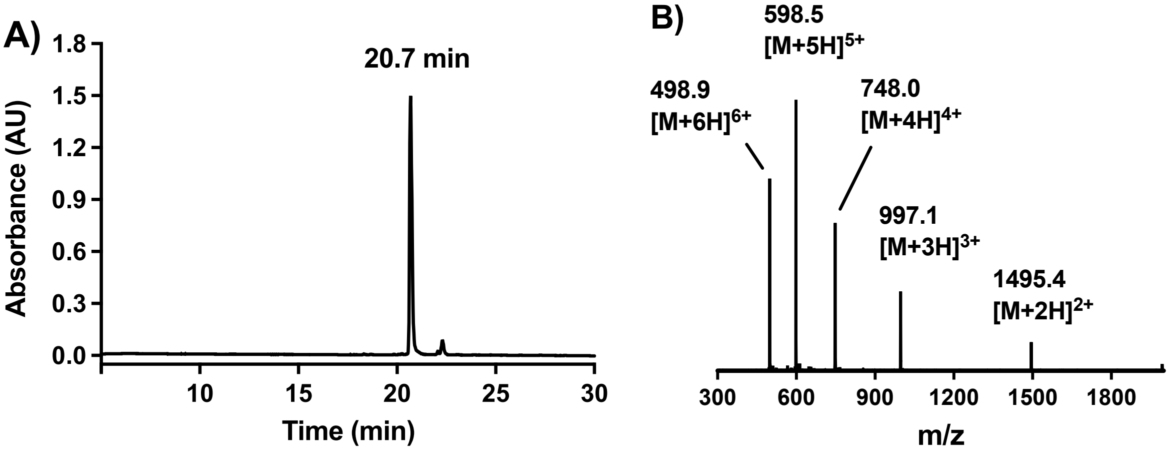


**Figure S5. A)** Analytical HPLC trace of RP-HPLC purified TS(56-81) (**4**), $R_t$ = 20.7 min (1 to 50% B over 30 min, 0.1 vol.% TFA, λ = 214 nm); **B)** ESI-MS of **4**, Expected Mass (ESI+): 1495.3 $[M+2H]^{2+}$, 997.2 $[M+3H]^{3+}$, 748.1 $[M+4H]^{4+}$, 598.7 $[M+5H]^{5+}$, 499.1 $[M+6H]^{6+}$. Mass Found (ESI+): 1495.4 $[M+2H]^{2+}$, 997.1 $[M+3H]^{3+}$, 748.0 $[M+4H]^{4+}$, 598.5 $[M+5H]^{5+}$, 498.9 $[M+6H]^{6+}$.

**One-pot Assembly of Native Thrombostasin (1):**

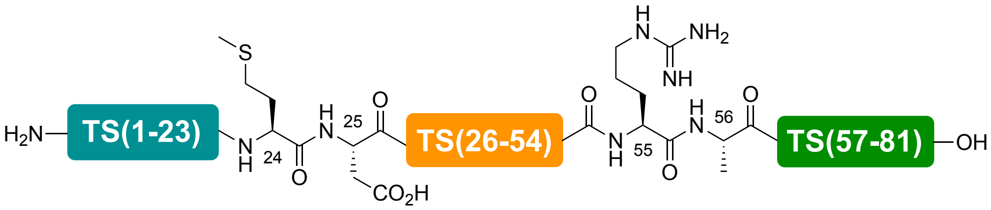


The N-terminal selenoester fragment **2** (1.4 mg, 0.52 µmol, 1.4 equiv. with respect to selenopeptide monomer) and bifunctional middle fragment **3** (1.5 mg, 0.38 µmol, 1 equiv.) were dissolved separately in ligation buffer (6 M Gnd•HCl, 100 mM $Na_2HPO_4$, pH 7.2) and combined together to carry out additive-free DSL (total reaction volume = 75 µL) as described in the general methods section. After completion of the DSL reaction in 10 min (based on UPLC-MS analysis), DPDS was extracted with 0.5 mL of $Et_2O$ (×5) and the pH of the reaction mixture was carefully was raised to 7.5 using 2 M NaOH for 5 minutes to hydrolyse the unproductive selenoester (resulting from trans-selenoesterification of excess N-terminal selenoester fragment by the selenol moiety of ligation product) and brought back down to pH 7.0. The ligation mixture was then treated with an equal volume of freshly prepared, degassed solution of 250 mM TCEP in ligation buffer (6 M Gnd•HCl, 100 mM $Na_2HPO_4$, pH 6.8, 50 equiv., 125 mM final concentration), resulting in complete deselenisation in 5 min as judged by UPLC-MS analysis.

To the crude ligation mixture, the cysteinyl fragment **4** (1.4 mg, 4.8 µmol, 1.3 equiv.) was added as a solid and the pH was adjusted to 6.8. After adding TFET (7.4 µL, 5 vol.%), the reaction mixture was incubated at 37 °C for 16 h to facilitate NCL. Finally, a solution of freshly prepared, degassed solution (150 µL) of 500 mM TCEP and 200 mM GSH in ligation buffer (6 M Gnd•HCl, 100 mM $Na_2HPO_4$, pH 6.8) was added to the reaction mixture, followed by VA-044 (2 mg, 20 mM final concentration). Incubation of the resulting mixture at 37 °C for 16 h followed by reverse-phase HPLC purification (0 to 20% B over 2 min then 20 to 80% B over 60 min, 0.1 vol.% TFA) afforded full-length native thrombostasin (**1**) as a white fluffy solid (0.7 mg, 20% over 4 steps).

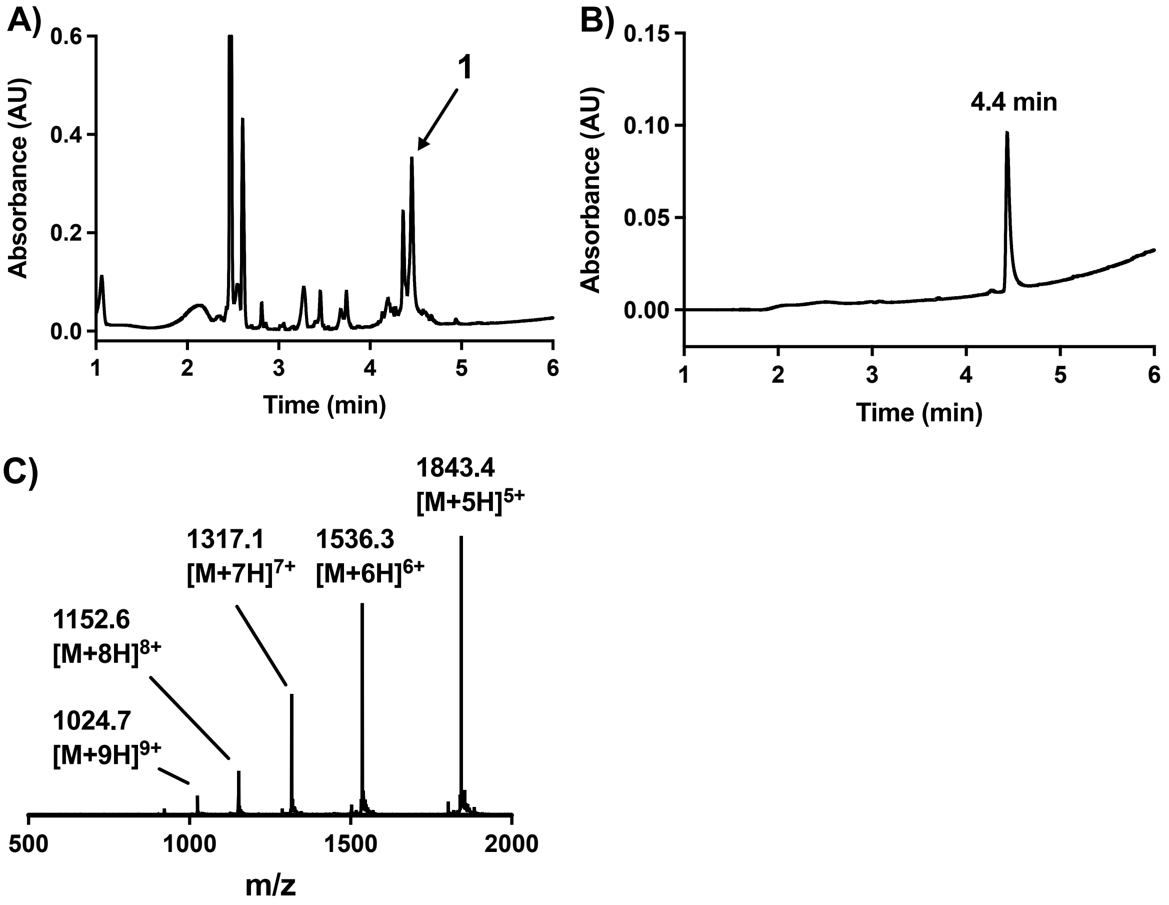

**Figure S6. A)** Analytical UPLC trace of crude reaction mixture after one-pot assembly of thrombostasin (**1**) via DSL-deselenisation and NCL-desulfurisation (gradient: 0 to 70% B over 5 min, 0.1 vol.% TFA, $\lambda$ = 230 nm). **B)** Analytical UPLC trace of RP-HPLC purified **1**, $R_t$ = 4.1 min (0 to 70% B over 5 min, 0.1 vol.% TFA); **C)** ESI-MS of **1**, Expected Mass (ESI+): 1843.3 $[M+5H]^{5+}$, 1536.2 $[M+6H]^{6+}$, 1316.9 $[M+7H]^{7+}$, 1152.4 $[M+8H]^{8+}$, 1024.5 $[M+9H]^{9+}$. Mass Found (ESI+): 1843.4 $[M+5H]^{5+}$, 1536.3 $[M+6H]^{6+}$, 1317.1 $[M+7H]^{7+}$, 1152.6 $[M+8H]^{8+}$, 1024.7 $[M+9H]^{9+}$.

**One-pot Assembly of Hepta-Glu Variant of Thrombostasin (7):**

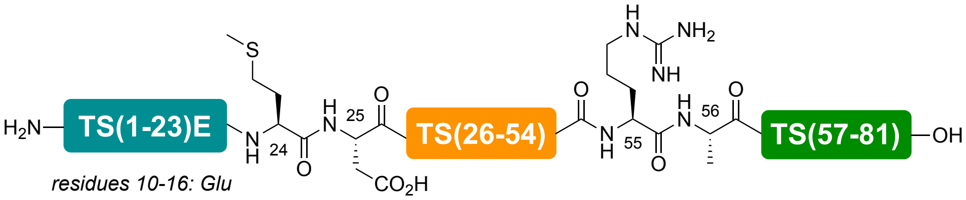


The TS(1-24)10-16E selenoester fragment **6** (8.8 mg, 3.12 µmol, 1.2 equiv. with respect to selenopeptide monomer) and bifunctional middle fragment **3** (10.2 mg, 2.61 µmol, 1 equiv.) were subjected to an additive-free DSL reaction (6 M Gnd•HCl, 100 mM $Na_2HPO_4$, pH 7.2; total reaction volume = 522 µL) using conditions described in the general methods section. After completion of the DSL reaction in 10 min (based on UPLC-MS analysis), DPDS was extracted with 0.5 mL of $Et_2O$ (×5) and the pH of the reaction mixture was carefully was raised to 7.5 using 2 M NaOH for 5 minutes to hydrolyse the unproductive selenoester (resulting from trans-selenoesterification of excess N-terminal selenoester fragment by the selenol moiety of ligation product) and brought back down to pH 7.0. The ligation mixture was then treated with an equal volume of freshly prepared, degassed solution of 500 mM TCEP in ligation buffer (6 M Gnd•HCl, 100 mM $Na_2HPO_4$, pH 6.8, 100 equiv., 250 mM final concentration), resulting in complete deselenisation in 5 min as judged by UPLC-MS analysis.

To the crude ligation mixture, cysteinyl fragment **4** (1.4 mg, 4.8 µmol, 1.3 equiv. with respect to selenopeptide monomer) was added as a solid and the pH was carefully adjusted to 6.8. Due to the unavailability of TFET, methyl thioglycolate (MTG) was used as an exogenous thiol additive in this case. After adding MTG (10.4 µL, 1 vol.%), the reaction mixture was incubated at 37 °C for 16 h to facilitate NCL. Finally, a solution of freshly prepared, degassed solution (1.04 mL) of 500 mM TCEP and 200 mM GSH in ligation buffer (6 M Gnd•HCl, 100 mM

$Na_2HPO_4$, pH 6.8) was added to the reaction mixture, followed by VA-044 (13.4 mg, 20 mM final concentration). Incubation of the resulting mixture at 37 °C for 16 h followed by reverse-phase HPLC purification (0 to 60% B over 65 min, 0.1 vol.% TFA) afforded full-length hepta-Glu variant of thrombostasin (**7**) as a white fluffy solid (8.7 mg, 36% over 4 steps).

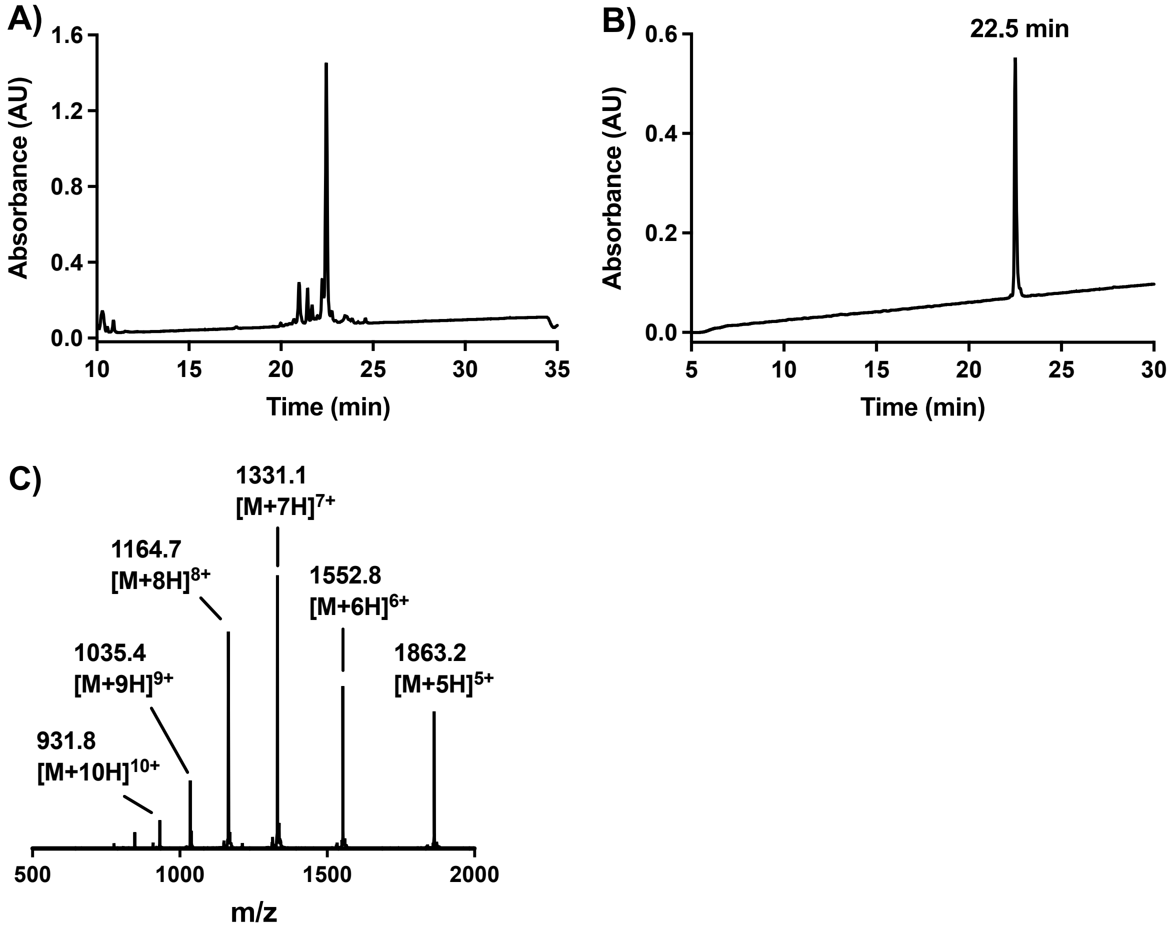


**Figure S7. A)** Analytical HPLC trace of crude reaction mixture after one-pot assembly of hepta-Glu variant of thrombostasin **7** via DSL-deselenisation and NCL-desulfurisation (gradient: 1 to 60% B over 30 min, 0.1 vol.% TFA, λ = 214 nm). **B)** Analytical HPLC trace of RP-HPLC purified **7**, $R_t$ = 22.5 min (1 to 60% B over 30 min, 0.1 vol.% TFA, λ = 214 nm); **C)** ESI-MS of **7**, Expected Mass (ESI+): 1863.2 $[M+5H]^{5+}$, 1552.9 $[M+6H]^{6+}$, 1331.2 $[M+7H]^{7+}$, 1164.9 $[M+8H]^{8+}$, 1035.6 $[M+9H]^{9+}$, 932.1 $[M+10H]^{10+}$. Mass Found (ESI+): 1863.2 $[M+5H]^{5+}$, 1552.8 $[M+6H]^{6+}$, 1331.1 $[M+7H]^{7+}$, 1164.7 $[M+8H]^{8+}$, 1035.4 $[M+9H]^{9+}$, 931.8 $[M+10H]^{10+}$.

**Synthesis of C-terminally Truncated Thrombostasin-Glu Variant (8):**

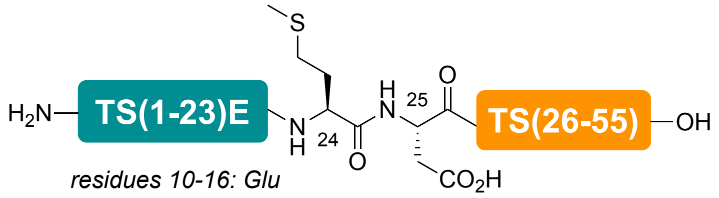


The TS(1-24)10-16E selenoester fragment **6** (2.6 mg, 0.92 µmol, 1.2 equiv. with respect to selenopeptide monomer) and bifunctional middle fragment **3** (3.0 mg, 0.77 µmol, 1 equiv.) were subjected to an additive-free DSL reaction (6 M Gnd•HCl, 100 mM $Na_2HPO_4$, pH 7.2; total reaction volume = 154 µL) using conditions described in the general methods section. After completion of the DSL reaction in 10 min (based on UPLC-MS analysis), DPDS was extracted with 0.5 mL of $Et_2O$ (×5) and the pH of the reaction mixture was carefully was raised to 7.5 using 2 M NaOH for 5 minutes to hydrolyse the unproductive selenoester (resulting from trans-selenoesterification of excess N-terminal selenoester fragment by the selenol moiety of ligation product). The ligation mixture was then treated with an equal volume of freshly prepared, degassed solution of 500 mM TCEP in ligation buffer (6 M Gnd•HCl, 100 mM $Na_2HPO_4$, pH 6.8, 100 equiv., 250 mM final concentration), resulting in complete deselenisation in 5 min as judged by UPLC-MS analysis. Finally, the thioester was hydrolysed to the corresponding carboxylic acid by incubating the reaction mixture at pH 8.0 for 8 h, followed by reverse-phase HPLC purification (0 to 60% B over 65 min, 0.1 vol.% TFA) to afford the C-terminally truncated hepta-Glu variant of thrombostasin (**8**) as a white fluffy solid (3.0 mg, 61% over 2 steps).

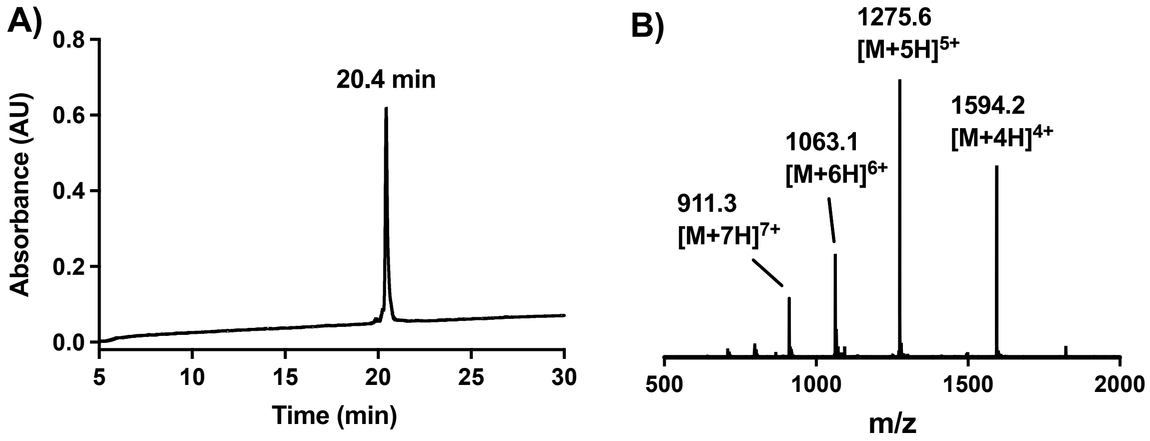


**Figure S8. A)** Analytical HPLC trace of RP-HPLC purified TS(1-55)E **8**, $R_t$ = 20.5 min (1 to 60% B over 30 min, 0.1 vol.% TFA, λ = 214 nm); **B)** ESI-MS of **8**, Expected Mass (ESI+):

1594.2 $[M+4H]^{4+}$, 1275.6 $[M+5H]^{5+}$, 1063.1 $[M+6H]^{6+}$, 911.4 $[M+7H]^{7+}$. Mass Found (ESI+): 1594.2 $[M+4H]^{4+}$, 1275.6 $[M+5H]^{5+}$, 1063.1 $[M+6H]^{6+}$, 911.3 $[M+7H]^{7+}$.

**Thrombin Inhibition Assay**

The inhibition of the amidolytic activities of human α-thrombin (Prolytix) by thrombostasin (**1**) and the hepta-Glu variant of thrombostasin and its C-terminally truncated analogue (**7** and **8**, respectively) was monitored spectrophotometrically using Tos-Gly-Pro-Arg-pNA (Chromozym TH, Roche) as chromogenic substrate. The Hepta-Glu variant of thrombostasin and its C-terminally truncated analogue (**7** and **8**) were also assessed against γ-thrombin using the same chromogenic substrate. The assays were performed in 50 mM Tris-HCl pH 8.0, 50 mM NaCl, 1 mg $mL^{-1}$ bovine serum albumin, with 0.2 nM human α- or γ-thrombin, 100 μM substrate, and varying concentrations of a given inhibitor. The concentration of each inhibitor was determined using an infrared spectrometer (Direct Detect, Millipore). All reactions were initiated by the addition of thrombin and were carried out at 37 °C in 96-well flat-bottom microtiter plates. Reaction progress was monitored at 405 nm for 40 min, with measurements taken every 5 min on a multi-mode microplate reader (Spark 10M, Tecan). All measurements were performed in duplicate. Log-dose response curves were generated by calculating the percentage of inhibition relative to the control in the absence of inhibitor, and $IC_{50}$ values were determined by fitting these curves to a nonlinear regression model using Prism 9 (GraphPad software).

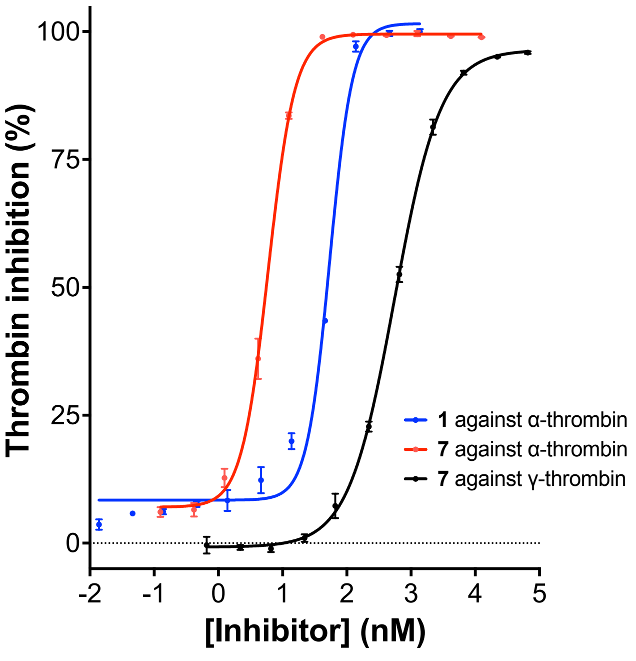

**Figure S9.** Dose-response curves for the inhibition of α- and γ-thrombin by thrombostasin analogues **1** and **7**. $IC_{50}$ of **1** against α-thrombin = 54 nM; $R^2$ = 0.9911; $IC_{50}$ of **7** against α-thrombin = 5.97 nM; $R^2$ = 0.9989; $IC_{50}$ of **7** against γ-thrombin = 556 nM; $R^2$ = 0.9994.

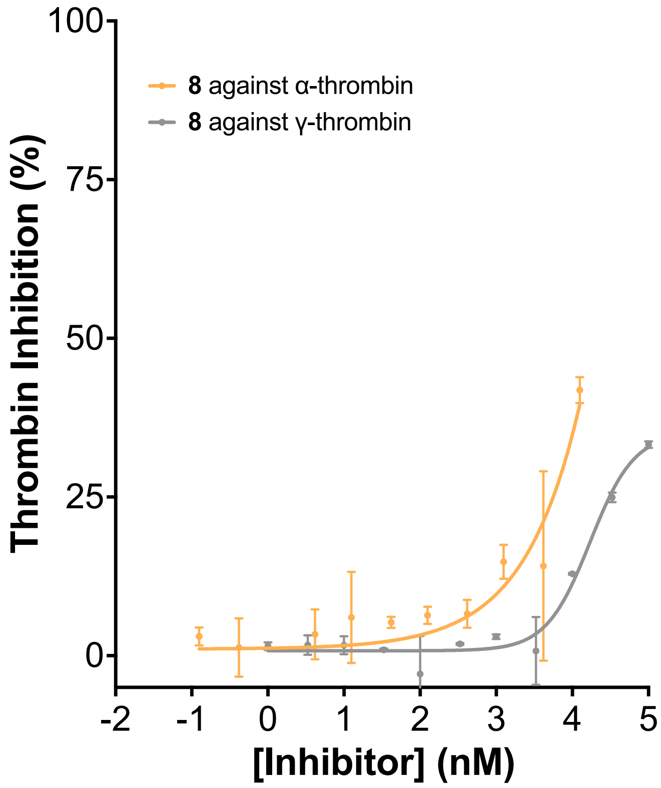


**Figure S10.** Dose-response curves for the inhibition of α- and γ-thrombin by C-terminally truncated hepta-Glu variant of thrombostasin **8**. $IC_{50}$ > 10 µM against both thrombin isoforms.

#### Enzymatic Cleavage of Thrombostasin

The hepta-Glu variant of thrombostasin **7** (25 µM) was incubated with α-thrombin (human, 1 µM, Prolytix) in assay buffer (25 mM Tris–HCl pH 8.0, 150 mM NaCl) at 37 °C. At the desired timepoints (t = 1, 5, 15, 60 min), the aliquots were quenched with aqueous 0.1 vol.% formic acid solution. Each timepoint was injected onto a Shimadzu 2020 LC-MS instrument with an LCM20A pump, SPD-20A UV/Vis detector and a Shimadzu 2020 (ESI) mass spectrometer running in positive mode utilising a Waters Sunfire 5 µm, 2.1 × 150 mm column (C18) at a flow rate of 0.6 mL $min^{-1}$.

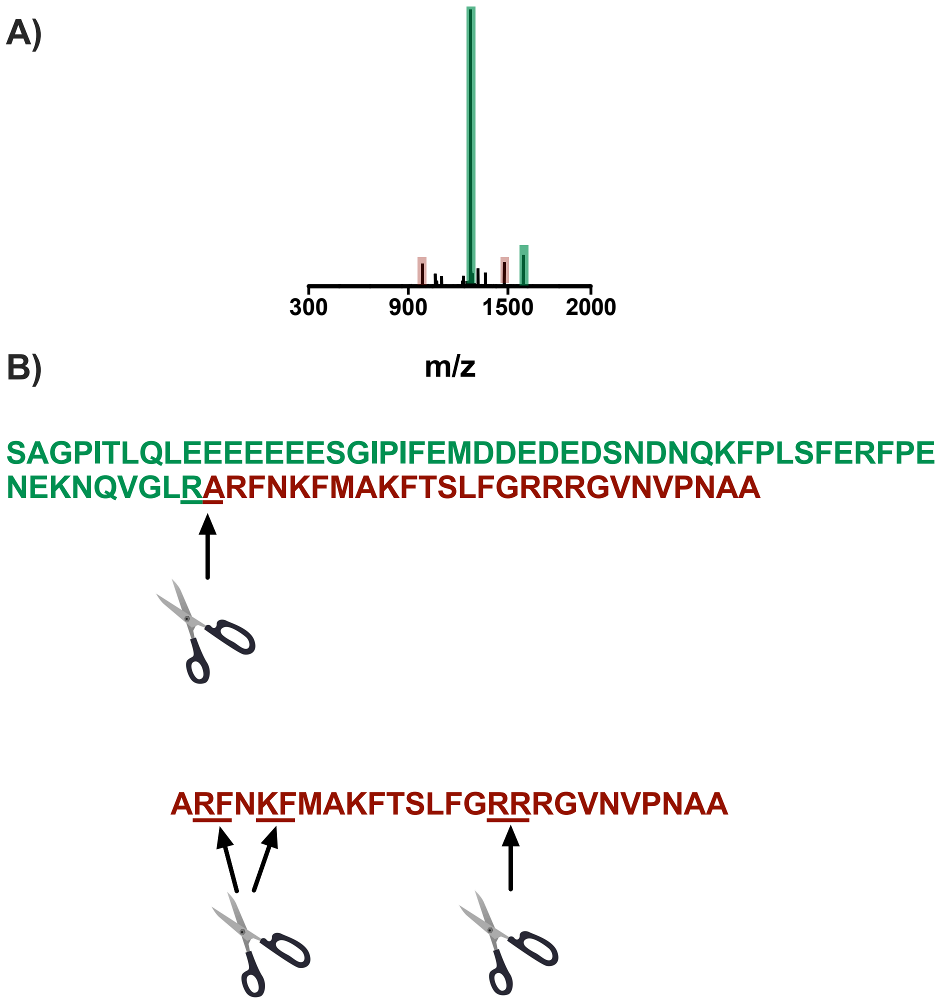


**Figure S11. A)** ESI-MS of **7** after 1 min incubation with α-thrombin; **B)** Amino-acid sequence of full-length **7** wherein underlined residues indicate cleavage sites.

##### activated Partial Thromboplastin Time (aPTT) assay

Activated partial thromboplastin time (aPTT) was measured using a BFT II analyser (Siemens Healthineers). Test compounds were preincubated with citrated platelet-poor human plasma (Diagnostica Stago) for 2 min at 37 °C. Plasma samples (50 µL) were then mixed with Actin FSL reagent (50 µL) and incubated for a further 180 s at 37 °C. Coagulation was initiated by addition of pre-warmed $CaCl_2$ (50 µL), and clotting times were determined automatically using the BFT II analyser. Compounds were evaluated by serial dilution while maintaining a constant final vehicle concentration. Measurements were performed in quadruplicate, and aPTT values are reported in seconds.
